# Proliferation of superficial neuromasts during lateral line development in the Round Goby, *Neogobius melanostomus*

**DOI:** 10.1101/386169

**Authors:** J. M. Dickson, J. A. Janssen

## Abstract

Members of the family Gobiidae have an unusual lateral line morphology in which some of the lateral line canal segments do not develop or enclose. This loss of lateral line canal segments is frequently accompanied by proliferation of superficial neuromasts. Although the proliferation of superficial neuromasts forms intricate patterns that have been used as a taxonomic tool to identify individual gobiid species, there has never been a detailed study that has documented the development of the lateral line system in gobies. The Round Goby, *Neogobius melanostomus,* is the focus of this study because the absence of the lateral line canal segments below the eye are accompanied by numerous transverse rows of superficial neuromasts. Our results suggest that the origin of some of these superficial neuromast lines could be the result of single presumptive canal neuromasts that have proliferated after canal enclosure is arrested. Many of the intricate patterns of neuromasts observed in gobiids develop from a simplified pattern of neuromast that is very similar among different species of gobies. The proliferation of superficial neuromasts has evolved several times in fish families such as the tetras, gobies, and sculpins, and may provide an adaptive advantage to ‘tune’ the lateral line system for different environments and prey types.

**SIGNIFICANCE STATEMENT:** Understanding the development of different lateral line morphologies can provide insights into how these morphologies have convergently evolved in many fish taxa. This is the first study to document the progression of the development of the reduced lateral line morphology. This study shows evidence that the developmental origins of orthogonal lines of superficial neuromasts posterior to the eye are not neomorphic lines, but in fact arise from precursor neuromasts that seem to be analogous to presumptive canal neuromasts.

## INTRODUCTION

Not only do fishes occupy a broad range of habitats, they are extremely diverse and comprise over 24,000 species (Helfman *et al.*, 1997). Both the environment and phylogenetic histories are thought to influence the morphology of fish sensory systems such as the lateral line system. The lateral line system is a mechanosensory system that is used to detect water movement (Denton and Gray, 1988) that is present in all cyclostomes, fishes, and aquatic amphibians (Yamada, 1973; Lannoo, 1987a; b; Braun and Northcutt, 1997; Northcutt, 1997). The morphology of the lateral line system can be quite diverse among fish taxa and amphibians (Webb, 1989; Northcutt, 1997). It is thought that variations in lateral line canal morphologies have evolved as a result of both environmental factors (especially in lentic species) and phylogenetic relationships (Webb, 1989; Northcutt, 1997; Webb and Northcutt, 1997; Fuiman *et al.*, 2004). There are a number of studies that have tried to place the lateral line morphology in a phylogenetic context (Webb, 1989; Webb and Northcutt, 1997), however the adult morphology alone is not enough to truly understand the evolution of the lateral line system.

Morphological changes can result from either changes in duration, rate, onset, or offset (heterochrony) or from changes in the source or patterning of ontogenetic trajectories (Northcutt, 1997). It is possible that even subtle ontogenetic changes in patterning either spatially (heterotopy) or temporally (heterochrony) during development can result in very different adult phenotypes (Hall, 1999). By examining the development of the lateral line system in fishes with unusual lateral line morphologies we may gain insight into how different lateral line morphologies convergently evolved in several fish taxa.

Heterochrony in particular has been suggested to play a role in different lateral line morphologies (Northcutt, 1997; Webb and Northcutt, 1997). Although most adult fishes have a combination of superficial neuromasts and canal neuromasts (which are enclosed in a lateral line canal) (Coombs *et al.*, 1988), both of these sensory organs are present on the surface of the epithelium early in development (Tarby and Webb, 2003; Webb and Shirey, 2003). Ontogenetically all neuromasts start in the epidermis above the basement membrane and are present at or slightly after hatching. It is only later on during development, usually late in the larval period (Fuiman *et al.*, 2004), that the presumptive canal neuromasts enclose in canals. Northcutt (1997) described the stages of lateral line development beginning from the initial formation of ectodermal placodes through the completion of canal enclosure. Northcutt (1997) proposed that in some taxa the absence of canals or canal segments may be the result of lateral line development that is arrested before canal enclosure occurs (Northcutt, 1997). This would mean that some superficial neuromasts are actually homologues of canal neuromasts, which are sometimes referred to as replacement neuromasts (Northcutt, 1997; Webb and Northcutt, 1997). This is thought to be the case with some lungfishes and an outgroup analysis suggests that the superficial neuromasts present in amphibians may also be homologous to canal neuromasts that failed to enclose (Northcutt, 1997).

Although complete canal enclosure is thought to be a synapomorphy for bony fishes, and most cartilaginous fishes including placoderms, chondrichthyes, and acanthodii, there are taxa within these groups where there is an apomorphic loss of canal segments. The loss of canal segments in cranial canals is usually referred to as ‘reduced canal’ morphology. There are a number of fishes that exhibit the reduced canal morphology (for a review refer to table in Coombs et al., 1988), thus this morphology has convergently involved in numerous fish taxa.

A prime example of a family with reduced cranial lateral line canal morphology is Gobiidae, (Korn and Bennett, 1975; Webb, 1989; Ahnelt *et al.*, 2000; Ahnelt and Duchkowitsch, 2001; Ahnelt *et al.*, 2004; Bertucci *et al.*, 2012; Langkau *et al.*, 2012). Gobies are an excellent model organism for evolutionary studies because they are very diverse (over 2000 species) and they occupy a number of different habitats (Ahnelt and Goschl, 2003). Even within a given geographic area gobies can occupy a number of different habitats ranging from exposed littoral surfaces to swiftly running streams. Not only do gobies have a reduced lateral line canal morphology, which means more of their neuromasts are embedded in the surface of the epidermis (superficial neuromasts) instead of being enclosed in canals (canal neuromasts), there is also a lot of species level variation in the patterns of superficial neuromasts present (Toshiaki J, 1971). In fact the diversity in the gobiid lateral line system is so great that it has been used as a taxonomic tool for identifying different species of goby (Sanzo, 1911; Iljin, 1930; Akihito, 1986; Miller, 1986).

Even within populations of the same goby species there can be differences in the lateral line system. Along the west coast of the United States northern populations of the tidewater goby, *Eucyclogoius newberryi,* have complete canals while more southern populations have greatly reduced canal types (Ahnelt *et al.*, 2004). Since there is so much variation present in their lateral line morphology in gobiids, we chose a gobiid species that is locally available, the round goby, as a starting point to follow the development of the lateral line system in a fish with the reduced lateral line canal morphology.

The round gobies have a reduced canal morphology in which several, but not all, of the cranial lateral line canals fail to enclose. We hypothesize that the development of canal segments that completely enclose will be similar to the development that has been reported in other fishes. By comparing the development of neuromasts where canal segments form with the development of neuromast where canal segments do not form, it will be possible to identify if and where differences in neuromast development occur. Since presumptive canal neuromasts are often present early in development, this would provide information on which superficial neuromasts are most likely canal neuromast homologues (canal segments containing neuromasts are replaced by lines of superficial neuromasts).

This study should provide the foundations for answering broader questions about the relationship, if any, between neuromast proliferation and canal formation. What happens when a canal segment is deleted? Do presumptive canal neuromasts that do not enclose in canal segments behave like superficial neuromasts? Many species with the reduced canal morphology show neuromast proliferation in areas where canal segments are absent, such as Cobitidae (Lekander, 1949), Gobiidae (Pezold, 1993; Ahnelt *et al.*, 2004), Lophiiformes, and Cyprinodontidformes (Coombs *et al.*, 1988). We hypothesize that in the round goby the superficial neuromasts in regions where canal segments are lacking are homologous to canal neuromasts. It is thought that canal neuromasts do not proliferate after being enclosed in canals, but superficial neuromast proliferation may still occur during the post-embryonic period (Sato, 1955; Peters, 1973; Puzdrowski, 1989). If proliferation is a key difference between canal and superficial neuromasts, can stranded canal homologues regain their ability to proliferate?

## MATERIALS AND METHODS

In order to identify developmentally important time points in goby lateral line development, we sampled Round gobies, *Neogobius melanostomus,* from nests collected from Lake Michigan (see 16a). We sampled specimens 6-74 mm SL. Lateral line development was assessed in live fish using DASPEI staining and fish previously prepped for SEM to determine when morphological changes occurred.

### Animals

SCUBA divers collected goby egg nests attached to rocks from Lake Michigan. Nests were transported back to the Great Lakes WATER Institute in coolers during the summer of 2011. After about two weeks, the larvae were moved into large conical tanks. And either fed zooplankton collected from Lake Michigan, daphnia, or cultured *Artemia* nauplii and adults. Larger juveniles (36-61mm SL) were collected using a seine net.

### DASPEI

Fish were incubated in a 0.01% DASPEI (2-(4-(dimethylamino)styryl) -N-ethylpyridinium iodide, Invitrogen, Carlsbad, CA) in Danieau buffer (trace salt solution buffered with HEPES (pH 7.6)) used to raise zebrafish embryos/larvae for 30 min. to an 1 hour. DASPEI is a vital stain that stains the mitochondria in the hair cells of the lateral line neuromasts. Gobies were then anesthetized in 0.1% Tricaine methanesulfonate (MS-222) in 1X Danieau buffer. Fish were placed on a Petri dish before they were viewed with a Leica MS75 microscope with a blue laser to excite the DASPEI staining. Images were taken using a Nikon coolpix 8800 camera attached to the ocular. After, the larvae were either fixed in 10% formalin or in Karnovsky’s fixative for SEM. DASPEI staining was used to monitor the development of the lateral line system, which facilitated a more precise sampling schedule.

### Scanning Election Microscopy

A size series of gobies ranging from 6 mm-38 mm SL were preserved in Karnovsky’s fixative (2% paraformaldehyde and 2.5% glutaraldehyde in 0.1M cacodylate buffer pH 7.4). They were kept at room temperature for several days to facilitate penetration of the fixative and afterwards stored at 4°C. Before they were dehydrated they were rinsed in distilled water. Fish larger than 14 mm SL were cut into sections before they were immersed in 1% osmium tetroxide for 1 hour (Janssen *et al.*, 1987) and dehydrated in a graded ethanol series (50%, 70%, 95%, 100%, 100%) for thirty min for each step (Ahnelt and Bohacek, 2004). Samples were placed in an ultrasonic bath for 2 min in 70% ethanol to remove the cupula (Maruska and Tricas, 2004) and critical point dried. They were mounted on an aluminum stub. Samples were then sputter coated with an Emitech K575X sputter coater with either 10 nm iridium or gold coated palladium. They were then viewed on a Hitachi S-4800 FE SEM using the Quartz PCI program. Plates were assembled in photoshop and any measurements taken were done in ImageJ.

### Lateral line system descriptions

For sake of consistency with other gobiid studies the nomenclature used to described the lateral line canals and superficial neuromast lines in the round goby was based on the previous descriptions of the gobiid lateral line system (Ahnelt and Duchkowitsch, 2001; Ahnelt and Goschl, 2003; Ahnelt *et al.*, 2004). This nomenclature may differ from the terms used to describe the lateral lines in other species. For instance, the oculoscapular canal which is referred to in gobiid literature is commonly described as several different canals (such as supraorbital, otic, preopercular canals) in other fish species. ANCOVA statistical analyses were carried out with either ‘R’ or JMP Pro 13.

## RESULTS

Neuromast identification is based on Ahnelt papers (Sanzo, 1911; Miller, 1986; Ahnelt et al., 2000; Ahnelt and Duchkowitsch, 2001; Ahnelt and Goschl, 2003; Ahnelt and Bohacek, 2004; Ahnelt *et al.*, 2004), Sanzo, and Miller et al. (Fig.1). There appears to be a proliferation of neuromasts in gobiids, especially in the regions where canal segments are missing (infraorbital, certain oculoscapular segments, and the supratemporal region). These lines of superficial neuromasts were divided into major regions such as the mandibular, otic, preopercular, and dorsal regions and are described below. These regions are similar to what has been described in adults in the goby literature, but they do not necessarily reflect the innervation or the developmental origins of specific neuromast lines.

We observed that the number of neuromasts in all the major regions of the round goby significantly increase with fish size during early embryonic development (Figs. 2-6). and is described in detail below:

**Fig. 1.**
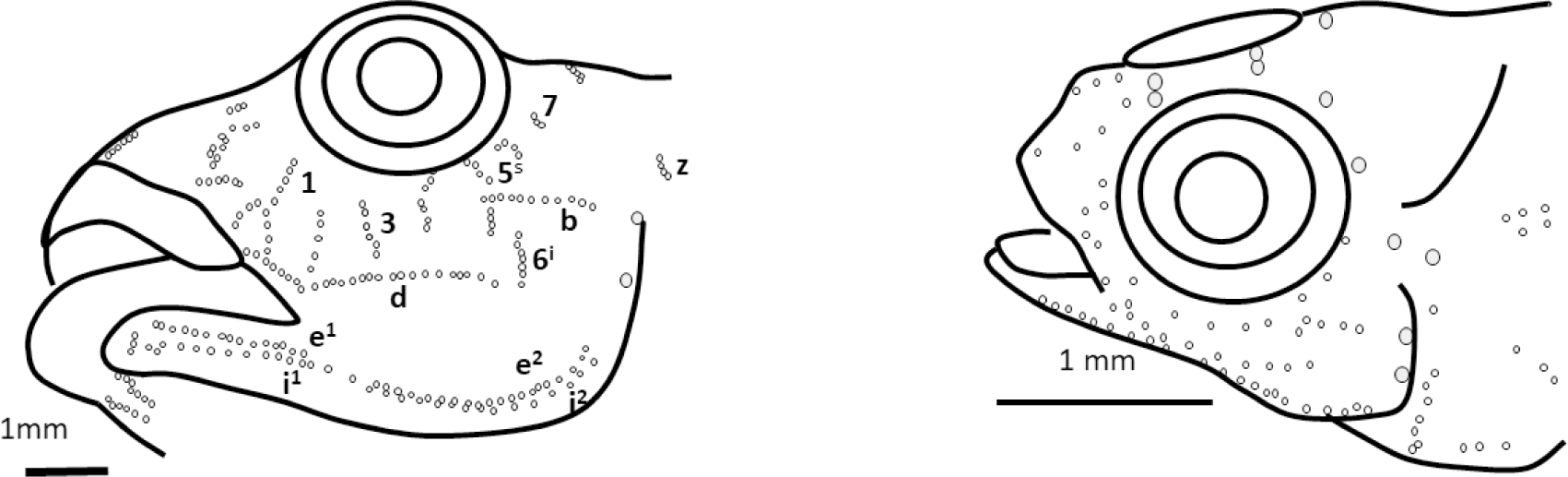
Diagram of neuromast position in Round goby, *Neogobius melanostomus*, at A) 30 mm SL and B) 9 mm SL. Small open circles indicate superficial neuromasts and larger open circles are canal neuromasts.

**Fig. 2.**
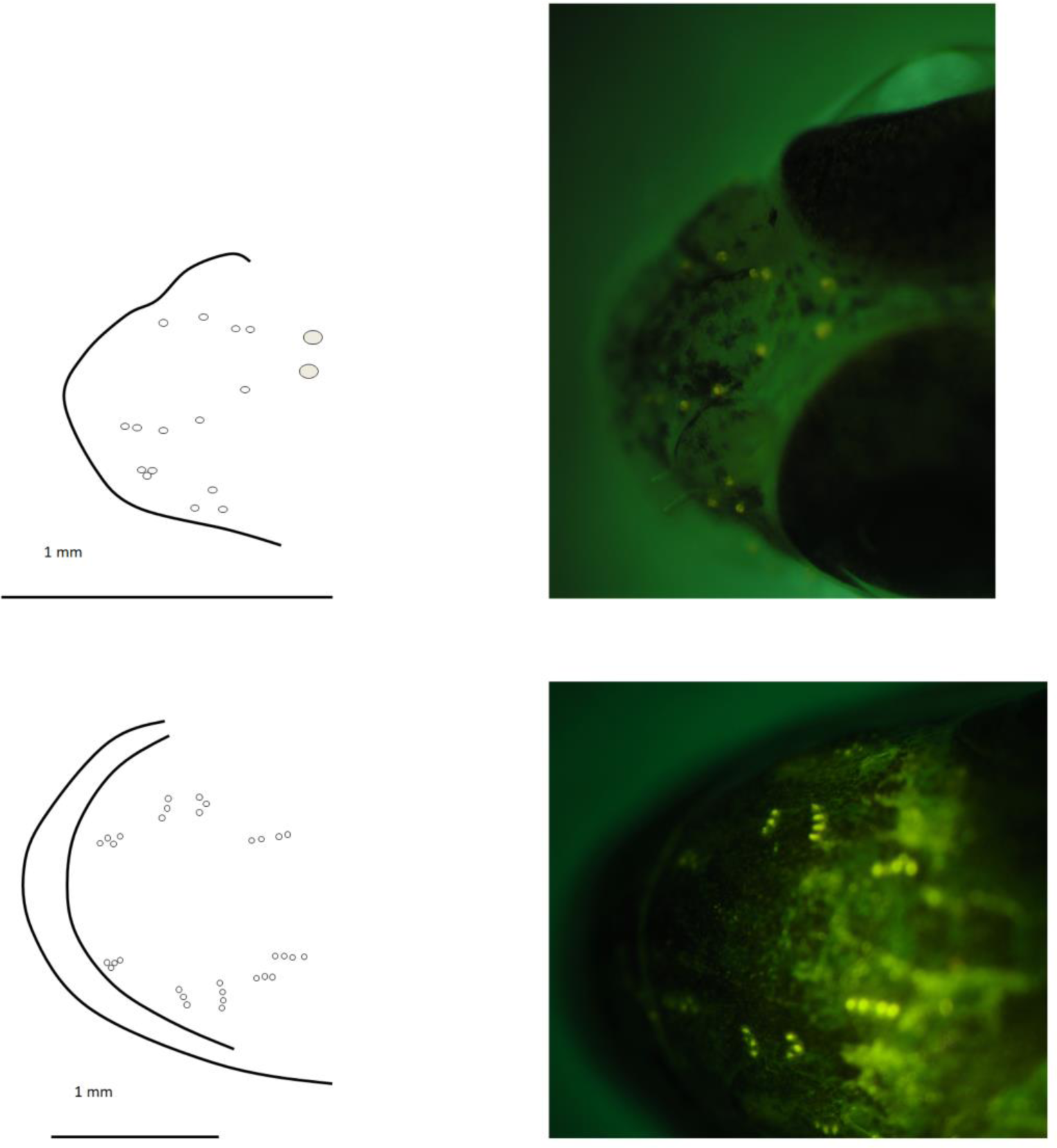
Dorsal view of Round Goby, *N. melanostomus*, heads throughout development at A) 12 mm SL, B) 30 mm SL. Neuromasts are fluorescently stained with DASPEI. Small open circles indicate superficial neuromasts and larger open circles are canal neuromasts.

### Preorbital (snout)

On the anterior portion of the head (snout), there are a median series lines (*r,* and *s* in other gobies). At standard length (SL) there were four neuromasts, and by 35 mm SL each of these four neuromast have proliferated into rows containing 3-4 neuromasts (Fig. 2). The two median rows are tangential, and the outer dorsal and ventral rows are longitudinal. The lateral series is comprised of indistinct rows belonging to *c*.

### Interorbital

Supraorbital canals are present, so it appears that there are no inter-orbital papillae, *p*, present between the eyes (Fig. 6).

**Fig. 6.**
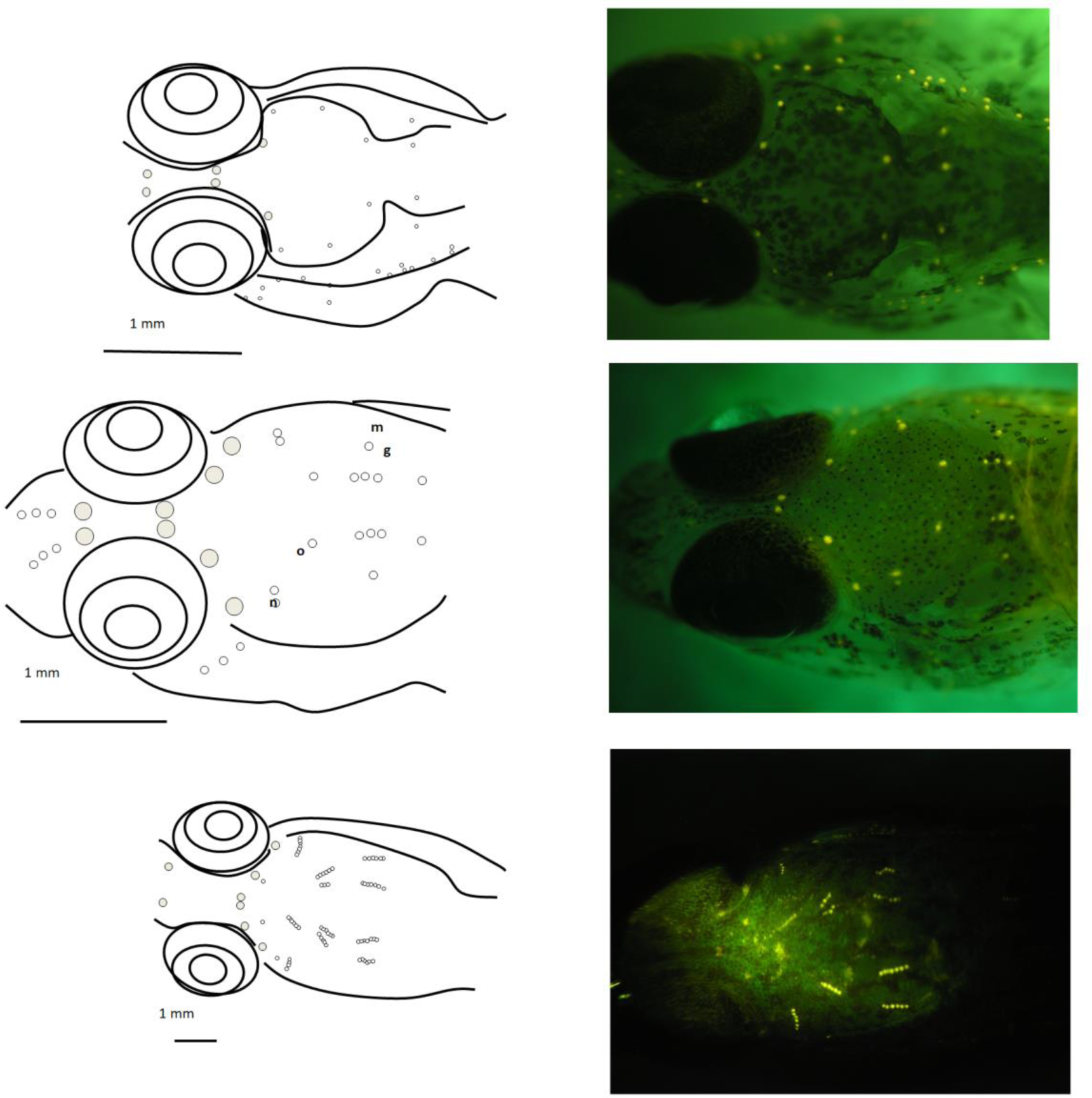
Dorsal view of Round Goby, *N. melanostomus*, heads throughout development. A) 9 mm SL, B) 12 mm SL C) 35 mm SL. Neuromasts are fluorescently stained with DASPEI. Small open circles indicate superficial neuromasts and larger open circles are canal neuromasts.

### Suborbital (infraorbital region below eye)

There are two rows of longitudinal neuromasts *b*, and *d* that increase in number throughout the early larval development. These rows are visible in the smallest fish examined 6 mm SL. At this time *b* contains ~2 neuromasts and *d*=11 neuromasts. In the largest fish examined 74 mm SL, there were 18-26 neuromasts in *b* and 66 neuromasts in *d*. There are typically 7 transverse (orthogonal) rows that form from neuromasts in the longitudinal row *a*. In a few individuals there is an extra row of neuromasts between lines *a4* and *a5* giving them a total of 8 transverse rows (ex: 58 mm SL individual). Two of these lines (*a5*, *a6*) are broken by the longitudinal line *b* and are thus divided into superior (*a5s, a6s*) and inferior (*a5i, a6i*). Line *a6i* can also be referred to as *cp*. Theses seven transverse lines are represented by a single neuromast in the smallest fish examined, 6 mm SL. At 11-12 mm SL neuromasts in these lines begin to proliferate and smaller neuromasts can be seen above and/or below the larger neuromasts that were initially present. There is an increase in the number of superficial neuromasts with fish size. By 74 mm SL there are up to 27 neuromasts in line *a1*, 12 neuromasts in line *a2*, 14 neuromasts in line *a3*, 10 neuromasts in line *a5s*, 5 neuromasts in line *a5i*, 6 neuromasts in line *a6s*, 12 neuromasts in line *a6i*, 3 neuromasts in line *a7* (Fig. 1, 3). ANCOVAs were run on the transverse lines of the subocular region indicate that there is a significant increase (p<0.5) in lines 1,4,5i,5s, 6i, and 6s (Fig. 5).

**Fig. 3.**
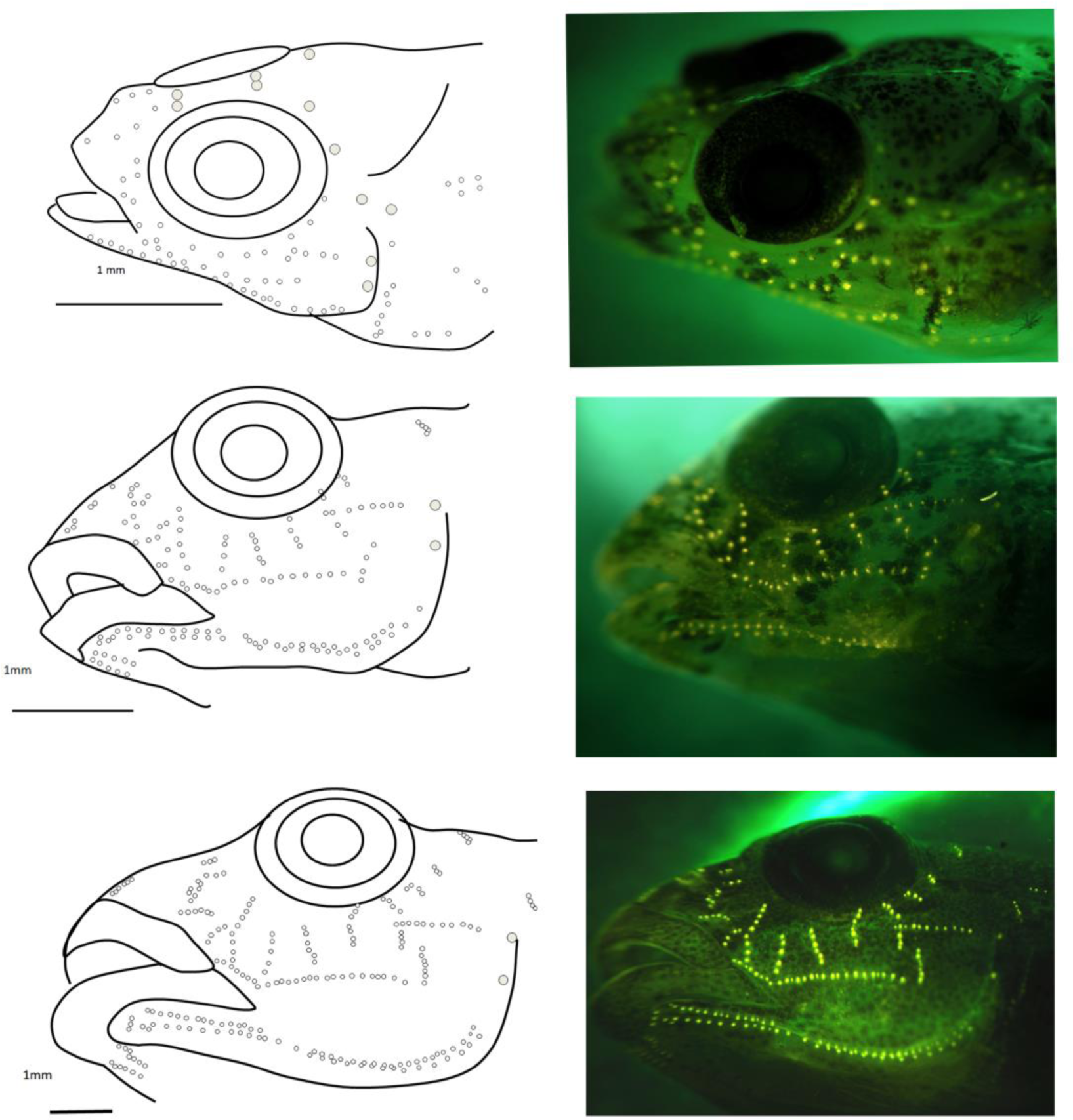
Lateral view of Round Goby, *N. melanostomus*, heads at different sizes during development. (File DSCN0909 A) is 9 mm SL, B) 16.5 mm SL(File DSCN1345) C) 30 mm SL. Neuromasts are fluorescently stained with DASPEI. Small open circles indicate superficial neuromasts and larger open circles are canal neuromasts..

**Fig. 5.**
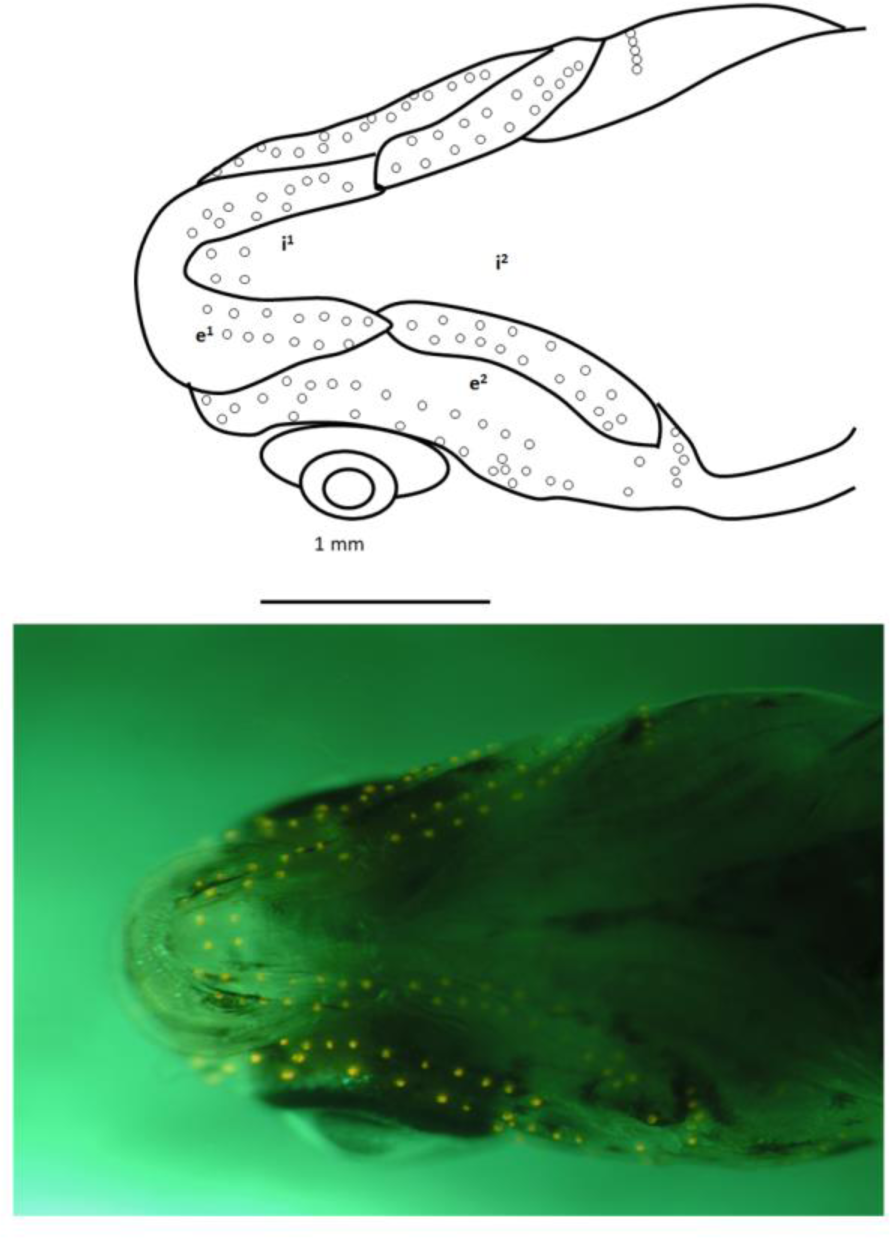
A) Ventral view of Round goby, *N. melanostomus*, at 9 mm SL. Neuromasts are fluorescently stained with DASPEI. Small open circles indicate superficial neuromasts and larger open circles are canal neuromasts

### Preopercular-mandibular

There are three longitudinal rows. Row *e* is on the lateral edge of the lower jaw and the preoperculum. This line is broken up into two lines, the mandibular line *e^1^* and the preopercular line *e^2^*. The medial row (*i*) is also broken up into a mandibular line, *i^1^*, and a preopercular line *i^2^*. At 6 mm SL there are 7 neuromasts in each of these rows, in larger individuals 35-75 mm SL there can be between 25-35 neuromast present in *i^1^* and up to 45 neuromasts in *i^2^*. There is also a medial row called *f*, which initially appears as a longitudinal line but by 38 mm SL *f* is more a triangular shaped cluster of neuromasts (Fig.3).

### Oculoscapular (Supraorbital/otic region)

There are transverse lines (*z*, *u^1^, u^2^*, *u^3^, u^4^*, *as^1-3^*) and longitudinal rows (*x^1^, x^2^, la^1-3^*). Row *z* is anterior and dorsal to the preopercular pore M. Initially, there are large neuromasts in what appears to be lines similar to longitudinal superficial neuromast lines *u*. These individual neuromasts proliferate to become three transverse lines ventral to *x^1^.* There is a second *x^2^* line that is posterior to canal segment K-L (Fig. 3).

### Opercular

There is a transverse row (*ot*) and two longitudinal rows, (*os, oi*). The *ot* has neuromasts ranging from 6 (6 mm SL) to 45 (74 mm SL). In the two longitudinal rows, *os* ranges from 2-21 (6-74 mm SL) whereas *oi* ranges from 4-18 (6-74 mm SL). In general, there does seem to be an increase in neuromast number within each row with size (Fig. 3).

### Anterior dorsal (occipital)

There are two transverse lines (*n, o*) and two longitudinal lines (*g, m, h*). The two transverse lines are more anterior than the other longitudinal rows. Row *n* is just posterior to the pore E, and *o* just anterior to the two longitudinal rows *g* (more medial) and *m* more lateral. Initially there is one neuromast present in each of these lines as early as 9 mm SL. By 74 mm SL there are up to ten neuromasts in the *n,* 13 in line *m,* and nine in row *g.* The superficial neuromast line *h* is located just posterior to the dorsal fin and is divided into two sections. Although *h* was only examined in four individuals, the number of neuromasts within these lines does seem to increase with age. At 14 mm SL there are three neuromasts in the anterior *h* line and four neuromasts in the posterior *h* line. The amount of neuromasts in these lines in a 35 mm SL individual was eight and six respectively (Fig. 6).

### Cranial canals

The pores for the supraorbital region of the oculoscapular canal B, C, D, E, F and pores F, G, H, K, L from the otic region are present on the oculoscapular. There is a fused pore C between the right and left canal segments. This means there are four canal segments in the supraorbital region, and three canal segments in the otic region. There are two canal segments in the preopercular region (M, N, O pores). By 5.5 mm SL there are grooves and ectodermal ridges beginning to form in the supraorbital and otic regions of the oculoscapular canal. By 6.5 mm SL some segments in the SO region of the oculoscapular canal are completely enclosed and the ectodermal ridges are present in the middle segment (G-H) of the otic region of the oculoscapular canal. There is some variation in the timing of canals enclosure, but by 27 mm SL all canal segments seem to be enclosed.

The anterior oculoscapular canal segments (in the supraorbital region) appears to enclose first (around 9 mm SL), followed by the posterior regions (enclosed by ~12 mm SL). The otic region has the anterior most segments enclosed by ~12 mm SL and posterior segments completely enclosed by 14 mm SL, and the preopercular region has one segment enclosed by ~12mm SL and complete enclosure can occur as early as 14.5 mm SL, however there were fish 16 mm SL that only had one preopercular segment enclosed. In the round goby complete canal enclosure occurs in many of the segments of the oculoscapular canal by 27 mm SL. In the preopercular region of the oculoscapular canal there is variation in which canal segments begin to enclose first. Most fish examined had the ventral region enclosing first, but there was one 14mm SL individual that had the dorsal region enclosed first. In the otic region of the oculoscapular canal it appears that the anterior segment (12 mm SL) encloses first followed by the anterior segment and then the posterior most segment (14.5 mm SL).

### Trunk

Individual neuromasts along the mid-line (*lm*) of the trunk are visible in SEM prepped individuals at 6 mm SL before scales have begun to form. Scales are present by 9 mm SL. By 10 mm SL clusters of two neuromasts are common along the midline of the trunk, and 22 mm SL there are clusters of 3-4 neuromasts, and by 38 mm SL there are clusters of 6 to 7 neuromasts on each scale long the midline. There is also a short line of neuromasts dorsal to the mid-line, *ld,* and another short line of neuromasts in the ventral region of the trunk, *lv*.

### Caudal fin

The caudal region was examined in 3 individuals with DASPEI staining. On the caudal fin there are three longitudinal lines of neuromasts: there is a dorsal line, *lcd*, the medial line, *lcm* and the ventral line, *lcv*. Although no statistical analyses were conducted due to the limited number of individuals sampled the smallest fish examined, 11 mm SL had 8 neuromasts in the dorsal most *lcd* line, 11 neuromasts in the medial line, *lcm*, and 13 neuromasts in the ventral *lcv* line. By 35 mm SL the number of neuromasts in these lines are 39, 42, 39 neuromasts respectively. There is a transverse *lct* line present on the caudal fin anterior to the three longitudinal lines.

## DISCUSSION

### Diversity in the adult morphology of lateral line system: Advantages

There are a wide range of lateral line system morphologies and specializations reported in fishes. Although there are lateral line system specializations that seem to be to associated with certain fish taxa, such as the lateral physic connection in clupeids (Coombs *et al.*, 1988), there can also be a wide range of specialization in the lateral line system of closely related species within the same family. These different lateral line morphologies can aid in ‘tuning’ the lateral line system depending on what frequencies are biologically relevant and how much hydrodynamic noise is in the environment (Janssen, 2003). Given that closely related species that live in different environmental conditions can have very different lateral line morphologies, it has been hypothesized that evolutionary origins of different lateral line morphologies could result from subtle changes during development of the lateral line system, such as the arrest of canal formation in species that lack a canal (Coombs *et al.*, 1988; Northcutt, 1997). Thus, identifying the similarities and differences in the development of the lateral line system may provide insight into the evolution of different lateral line morphologies.

Since different lateral line morphologies are thought to be specialized for picking up biologically relevant signals in a particular environment, it is not surprising that a closely related species within the same family that live in different habitats can have a wide range of lateral line morphologies. In gobies there tends to be a proliferation of superficial neuromasts, which have advantages and tradeoffs depending on the environments. While superficial neuromasts may be able to detect a broad range of signals, these signals include hydrodynamic signals form the environment and self-generated hydrodynamic signals that are created during swimming. If these fish inhabit hydrodynamically noisy environments where there is a lot of water flow, or they are highly mobile species, superficial neuromasts may not be able to detect other biological signals that are relevant. Thus, having a particular combination of both superficial neuromasts and canal neuromasts may be beneficial in certain environments but not others, which may be why there is such a diversity of lateral line system morphologies in gobies and other fishes.

Even if there is a proliferation of neuromasts there may be some advantage to retaining some canal segments. Superficial neuromasts can detect a broad range of hydrodynamic signals, whereas neuromasts enclosed in lateral line canals can filter out the lower frequency environmental ‘noise’ such as turbulence due to the mechanical properties of the canal (Janssen, 2003). Since there is more resistance to water movement because of the presence of the canal walls, there is less water movement in the canals at lower frequencies, which means the canals can act as high-pass filters to block out low frequency noise (Janssen, 2003). At higher frequencies there is less impedance in the canal and the boundary layers take longer to form, thus canals are better at detecting high frequency signals such as those generated by prey, when there is a lot of low frequency background noise in the environment (Janssen, 2003). This allows for fishes in environments with low frequency noise to detect higher frequency signals such as those generated by prey, thus fishes can have different combinations of superficial and canal neuromasts to detect signals in their environment.

Loss of canal segments and the proliferation of superficial neuromasts is common in gobiids as well as other fish taxonomic groups such as the sculpins of Lake Biakal. It is also a common adaptation for fishes that live in caves, such as the Mexican cavefish, as well as those that live in tubid waters such as pirate perch, *Aphredoderus sayanus,* and many deep-sea fishes. Even within a single taxonomic group there can be drastic modifications in the lateral line system. In the sculpin family Abyssocottidae of Lake Baikal there can be a complete absence of canal segments, that have been replaced with superficial neuromasts on papillae (Sideleva, 1982; Janssen, 2003). In other cases, the entire canal is not lost, but canal segments are missing, which has been documented in both sculpins and gobies. One example of this occuring in the sculpins is the Anadyr River form of *Cottus cognatus*, which is missing some of the canal segments that are present in the Lake Michigan population (Sideleva, 1982). The presence or absence of different canal segments within the same species present in different environments also occurs in Gobiids. The Tidewater goby, *Eucyclogobius newberyi,* has latitudinal variation in the supraorbital canal morphology with the northern populations with the full lateral line canal system while the southern population has more reduced canals were segments are actually missing (Ahnelt *et al.*, 2004). Given this information it is likely that canal loss and proliferation of superficial neuromasts present in the adult morphologies could be the result of changes or modifications that occur during development.

### Canal diversity in gobies

In gobies it is common for neuromast proliferation of superficial neuromasts to occur in places were canal segments or entire canals are missing, thus there seems to be a relationship between canal segment loss and neuromast proliferation. The loss of canal segments or entire canals is so common among different species of gobies and can be used as a character for taxonomic classification. In fact, characteristics of the oculoscapular (which can also be referred to as the supraorbital and otic canals) canal have been used to propose that subfamily Gobiinae is monophyletic (Pezold 2007). Canal segment loss and the proliferation of neuromasts has also been noted in the goby literature. Miller indicated that members of the family Gobiidae tend to lack a developed canal in the suborbital (also called the infraorbital canal) and supratemporal regions and have numerous superficial neuromasts (sensory papillae).

This reduction in lateral line canals and segments that is present in the Round Goby is consistent with other members of the family Gobiidae and subfamily Gobiinae. The Round goby completely lacks the infraorbital canal under the eye, and in other canals there are missing canal segments. Like many other members of the subfamily Gobiinae, the first pore (A) and the first canal segment in the oculoscapular canal is absent, and there appears to be a shared pore for pores C and D. Unlike other members of subfamily Gobiinae, the Round Goby retains the N-O segment in the preopercular region (Cooper *et al.*, 2007). Since we know that development of canal segments can be asynchronous in other species (Tarby and Webb, 2003; Bird and Webb, 2014; Webb *et al.*, 2014), it is possible that temporal changes during canal development in gobies can lead to the loss of canal segments.

At least some of the canals in the round goby appear to complete all the stages of canal development described in by others (Northcutt, 1997; Tarby and Webb, 2003; Webb and Shirey, 2003). In the round goby, it appears that some of the primary neuromasts are present in the smallest individual examined 4.5 mm SL. As with other fishes, the development of the canal segments seems to be asynchronous, which later fuse leaving a pore between segments. In the round goby there is one neuromast per canal segment which is typical of Actinopterygians and a few Sarcopterygians such as *Neoceratodus* and *Latimeria* (Webb and Northcutt, 1997). Understanding the lateral line system development may not only provide insight into potential mechanisms for how the intricate patterns of superficial neuromasts could have evolved among different gobiid species, it could also enhance our understanding of how variations in lateral line canal morphologies convergently evolved many times in numerous fish families. Perhaps there are subtle changes in the timing of development that can lead to the different lateral line canal morphologies observed in adult fishes.

### Variations in superficial neuromasts

In gobies canal loss along the head and trunk is often accompanied by the presence of superficial neuromast proliferation in the location where the canal would be present. This is the most prominent in the in the anterior lateral line system. Since superficial neuromasts on the head are the most sensitive, because the boundary layer is the thinnest due to the negative pressure gradient (Janssen, 2003; Windsor and McHenry, 2009), an increase in the number of superficial neuromasts may be beneficial in certain habitats. The most striking example of neuromast proliferation in the Round Goby is in the suborbital region below the eye, which can be arranged in different patterns.

These patterns of superficial neuromasts can range from the more ordered rows and lines, to merely aggregations on the head (Ahnelt and Bohacek, 2004). There are two common arrangements of superficial neuromasts in the infraorbital region. Some gobies have the ‘transverse’ pattern (vertical or orthogonal lines of superficial neuromasts), while others have the longitudinal (horizontal) lines of neuromasts (Korn and Bennett, 1975). In general *N. melanostomus* seems to have the transverse pattern of neuromasts in the infraorbital regions similar to the rock goby *Gobius paganellus* (Ahnelt 2001).

There is also canal loss accompanied with superficial neuromast proliferation in the dorsal view of the goby. There is no supratemporal canal, however there are larger neuromasts that proliferate and form lines *g* and *m*. The supraorbital canal is present above the eyes and along the anterior edges of the neurocranium. There are also superficial neuromasts that seem to continue along the lateral edges of the neurocranium in the posterior region of the neurocranium where the supraorbital canal is no longer present.

In the family Gobiidae the body does not have a trunk lateral line canal or modified scales, there are only exposed superficial neuromasts on the trunk. The most common pattern of superficial neuromasts on the caudal fin is three longitudinal rows (Ahnelt and Duchkowitsch, 2001). This plesiomorphic caudal neuromast pattern is present on the caudal region of the Round Goby and is similar to that of *Gobius niger*, unlike the more derived patterns with 4-8 longitudinal lines of neuromasts found in deep water gobies such as *Deltentosteus quadrimaculatus, Deltentosteus collonianus,* and *Aphia minuta* (Ahnelt and Duchkowitsch, 2001). Although all these lines of superficial neuromast were labeled according to the literature and followed throughout development, these names still need to be confirmed by looking at the innervation (Kornis *et al.*, 2012).

The pattern of the superficial neuromasts has been successfully used as a major characteristic for identifying individual species in the field (Pennuto *et al.*; Albert I, 1965). This is useful since the pattern of superficial neuromasts in adults does not change, whereas other features such as coloration may be more variable and fade in preserved specimens. The lateral line system seems to be a good diagnostic characteristic for identifying adults, (Toshiaki J, 1971) but it maybe a less than reliable characteristic for identifying larvae. Although these patterns have been documented in detail in adults, this is the first study to describe the ontogeny of the superficial neuromasts patterns throughout development.

### Superficial neuromasts throughout ontogeny

We found that a few superficial neuromasts in the round goby are indeed present at hatching (~ 4.5-6 mm SL), however the intricate patterns of superficial neuromasts that are found in adults are not present at hatching. The longitudinal lines in the infraobital region have several neuromast early in development, while the transverse lines only contain a single neuromast that do not begin proliferating till about 10-12 mm SL. The number of superficial neuromast in many of the superficial neuromast lines seems to still be increasing in largest fish, 74 mm SL fish. These findings are consistent with what is known about the development of the more well studied posterior lateral lines system in fishes; the early development is very simplified and increases in complexity later (Pichon and Ghysen, 2004).

### Development and evolution of superficial neuromasts

Similar to other fishes ((Jones and Janssen, 1992; Northcutt, 1997; Tarby and Webb, 2003), it seems that the superficial neuromasts are ontogenetic precursors of all canal neuromasts in the Round Goby (Webb and Northcutt, 1997). If this is the case, presumptive canal neuromasts that are stranded, could exhibit the properties of superficial neuromasts, such as neuromast proliferation to form ‘stiches’ which is present in the round goby.

There are a number of superficial neuromasts that could be potential canal neuromast homologues, which result from changes in the timing of canal development (i.e. canal development is halted before the neuromasts completely enclose in canals or before the canal begins to form). We hypothesize that the developmental origins of these single neuromasts in the suborbital region are actually presumptive infraorbital canal neuromasts that are stranded due to the lack of canal formation (Figs 3). We further hypothesize that once these canal neuromasts are stranded they can begin to exhibit the proliferation behavior observed in superficial neuromasts, which leads to the formation of the transverse lines of superficial neuromasts present in the suborbital region of adult Round Gobies.

There is further evidence that some of these superficial neuromasts are actually canal homologues. The three major lines of evidence that some of the superficial neuromasts originated from stranded canal neuromasts are 1) in some cases there is still the presence of a groove, indicating canal development was arrested before the canals were completely enclosed and thus by definition are superficial neuromasts, 2) the timing of the first appearance of the parent neuromasts that will give rise to these lines of superficial neuromasts, 3) the size of the neuromast when it first arises relative to other canal neuromasts. The mandibular and anterior portion of the preopercular canal of the Round Goby have all three of these characteristics. There are also neuromasts that seem to have a different ontogenetic origin. These ‘neomorphic’ neuromasts tend to arise later in development and are smaller in size relative to the canal neuromasts.

Not all of the stranded canal neuromasts show signs of any canal formation, however, they show the same developmental pattern of large primary neuromasts that appear at the same time as canal neuromasts and then later undergo proliferating by budding, forming lines with secondary neuromasts. Some of the neuromasts in the otic regions (line *u*) may be an example of canal neuromasts that are proliferating. There may also be stranded canal neuromasts in the dorsal region of the Round goby. In many fishes there is a supratemporal commissure that connects the two main body canals (Coombs et al., 1988). Although this canal is not present in the Round goby there are single primary neuromasts in the region that are present at 9 mm SL that could be homologous to the primary neuromasts. These neuromasts then proliferate and give rise to the lines *m* and *g* (Fig. 6). Most superficial neuromasts are either accessory lines or replacement lines for canals lost in evolution (Coombs et al., 1988).

### Neomorphic lines

Neomorphic lines of superficial neuromasts are accessory or independent of existing canals (Lekander, 1949; Disler, 1960). It is possible that other mechanisms may have formed these lines. There is no clear link to canal neuromasts in that area. Some of the otic lines (*x, la*) and opercular lines (*ot, oi, os*) suborbital (infraorbital region) the longitudinal lines *d, b* in the suborbital (infraorbital) region are neomorphic. In zebrafish, it has been shown that neural induction is involved in stich formation (Wada *et al.*, 2013a; Wada *et al.*, 2013b).

In the future, the mechanisms for neuromast proliferation in the gobies should be investigated in more detail. Although the mechanisms for neuromasts development and proliferation have been known to vary in posterior lateral line development across fish taxa (Pichon and Ghysen, 2004), neuromast proliferation and development have never been compared in closely related species such as gobies with diverse patterns of superficial neuromasts. It is thought that variation in the lateral line system is due to environmental and phylogenetic differences. Perhaps the diverse lateral line patterns found in gobies are the result of both their phylogenetic history and adaptations to the many diverse environments that gobies inhabit.

In fishes, it appears that there are potentially several sources from which the lines of superficial neuromasts could originate. Neuromasts can ‘bud’ from existing neuromasts from the primary neuromasts which has been described in the anterior lateral line system (Zipser and Bennett, 1973; Harding et al., 1981; Stone, 1933; Stone, 1937; Northcutt et al., 1994; Jones and Corwin, 1996; Mackenzie, 2012) or neomorphic lines can result from neural induction. In the Round Goby it appears that the ‘budding’ of presumptive canal neuromasts (primary neuromasts) could be a possible developmental origin for some of the transverse (orthogonal) lines of superficial neuromasts, particularly in the suborbital region below the eye. Thus, understanding lateral line development may be important for determining how diverse and complex lateral line patterns of superficial neuromasts arise from very simplified lateral line patterns in larval fishes.

This study shows that many of the intricate lines of neuromasts present in adults are not present or are greatly simplified in early larvae which is similar to other fishes (Pichon and Ghysen, 2004). The Round goby showed dramatic increases in the number of neuromasts present during the larval period (Fig. 2-6) and the patterns of superficial neuromasts change throughout development. This means that the lateral line system may not be a good diagnostic characteristic for differentiating an individual species of goby larvae from other species of goby larvae. It is possible that at hatching many species of gobies that have completely different patterns of superficial neuromasts as adults have very similar patterns of superficial neuromasts as larvae. Changes in the number of superficial neuromasts during development have taxonomical, ecological, and evolutionary implications.

## ACKNOWLEDGEMENTS

I would like to thank the members of the Udvadia and Janssen labs at the University of Wisconsin-Milwaukee for their help with collecting and maintaining fish in the laboratory. I would also like to thank Drs. Udvadia and Svoboda for the use of their microscopes, supplies, and many fruitful discussions on imaging and fish development. We would also like to thank H.A. Owen for help with the SEM preparation. This project was funded by a *Grant-In-Aid of Research from the National Academy of Sciences, administered by Sigma Xi, The Scientific Research Society.* This animal care protocol for this project (11-12 #2) was approved Institutional Animal Care and Use Committee at the University of Wisconsin-Milwaukee.

## CONTRIBUTIONS

The intellectual foundations of this project, i.e. reduced lateral line canal development was from J.A. Janssen. Fish were collected and housed by J.A. Janssen. Microscopy, data collection, and statistical analyses were conducted by J.M. Dickson. Manuscript draft preparation was done by J.M. Dickson while J.A. Janssen assisted in manuscript edits and intellectual guidance.

Appendices:

---

**Fig. 4.**
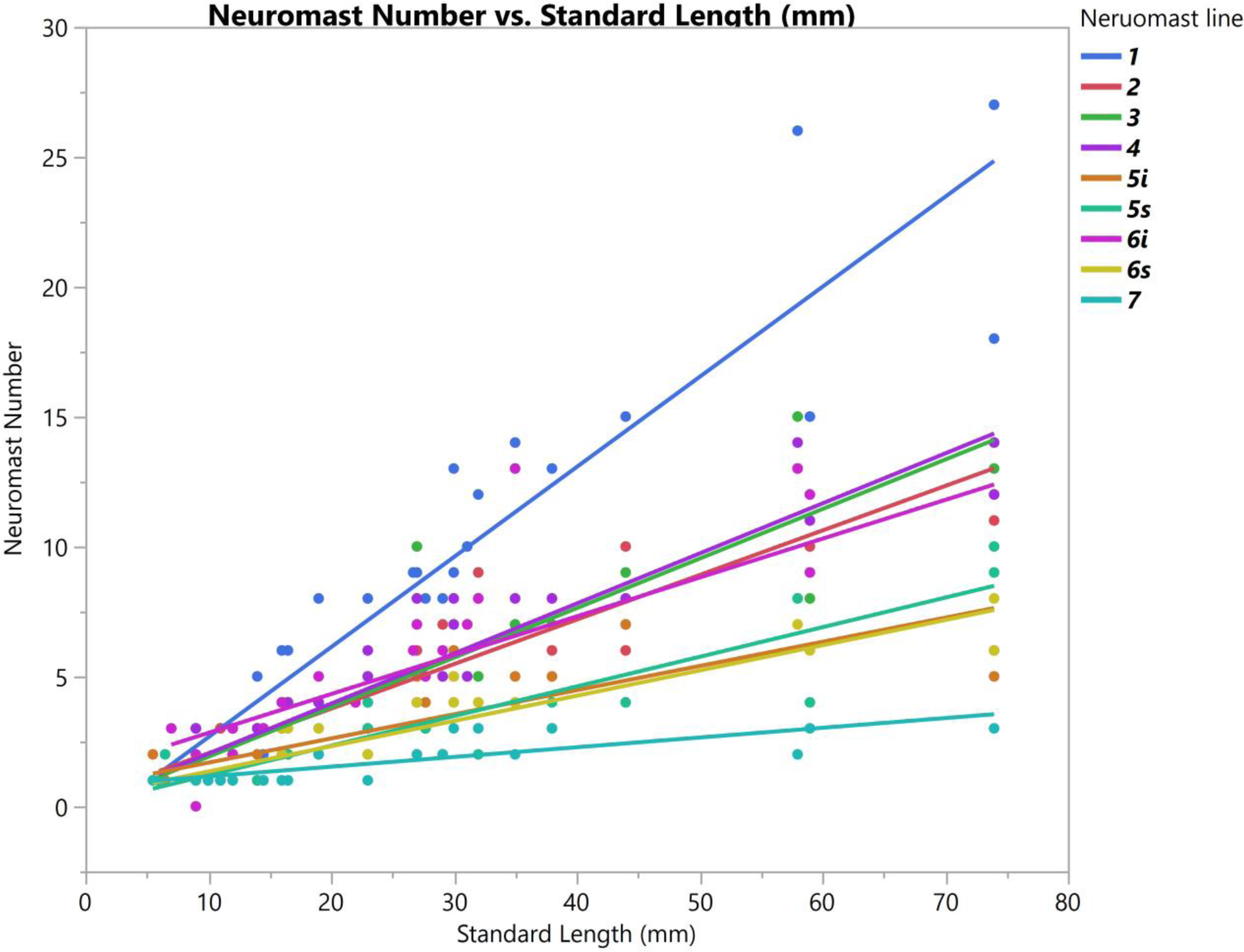
Number of superficial neuromasts in A) infraorbital region in superficial neuromast lines *1* (blue filled circles), *2* (red filled circles), *3* (light green filled circles), *4* (purple filled circles), *5s* (orange filled circles), *5i* (dark green circles), *6s* (gold filled circles), *a6i* (magenta filled circles), and *7* (light blue filled circles).

**Fig. 7.**
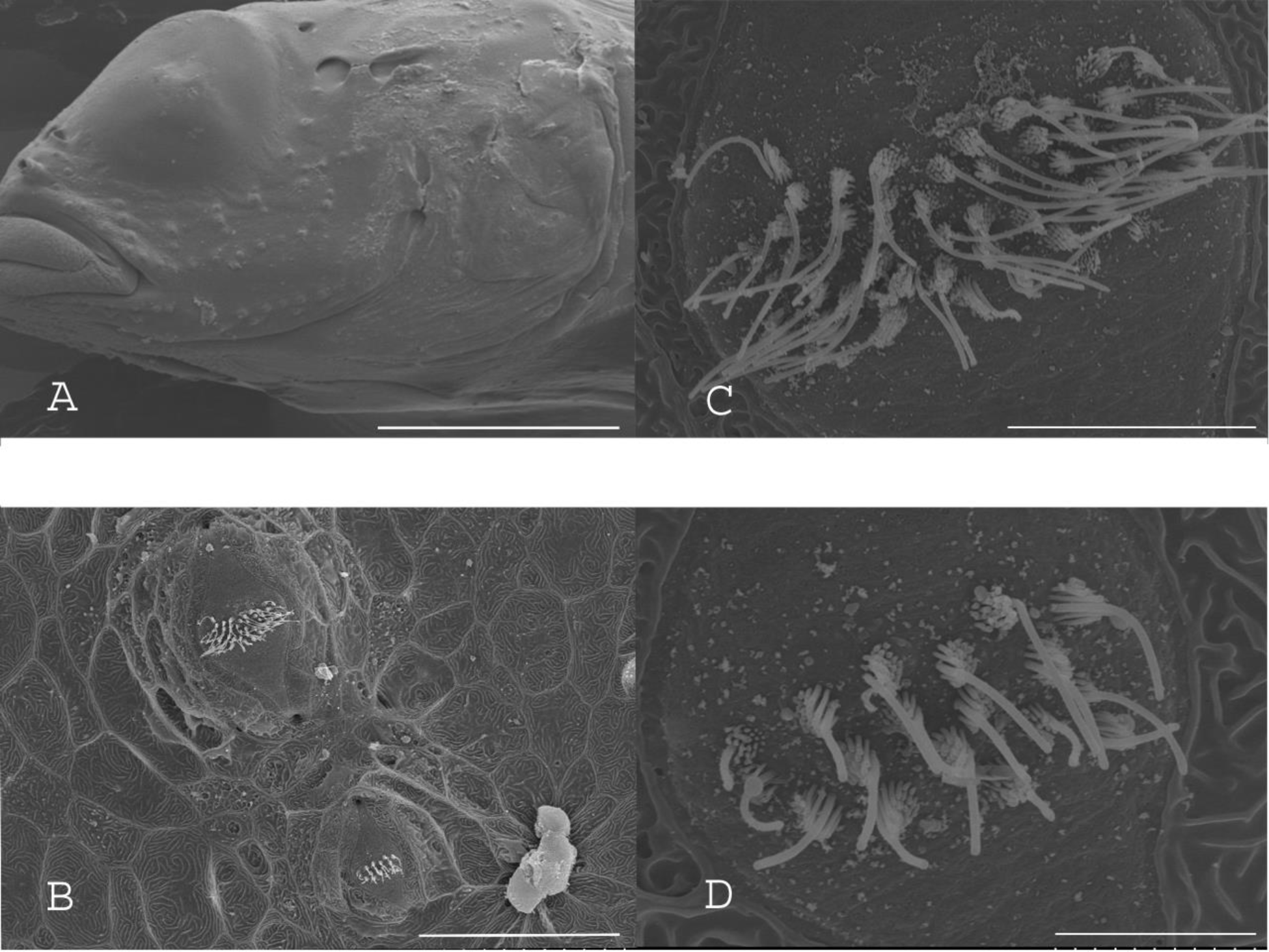
SEM micrograms of the superficial lines ventral to the eye in a 12 mm SL Round Goby, *N. melanostomus.* A) Lateral view of head taken at magnification of x45, B) proliferating neuromasts in the superficial lines of neuromasts below the eye at x800 magnification. C) The larger more dorsal neuromast pictured in ‘B’ at a higher magnification of x5000 and D) the smaller ventral neuromast pictured in ‘B’ magnified by x8000.

## Footnotes

1 Department of Biological Sciences, University of Wisconsin-Milwaukee, 3209 N. Maryland Ave, Milwaukee, WI, 53201.

2 School of Freshwater Sciences, University of Wisconsin-Milwaukee, 600 East Greenfield Ave, Milwaukee, WI, 53204.

